# 3D in-situ characterization reveals the instability-induced auxetic behavior of collagen scaffolds for tissue engineering

**DOI:** 10.1101/2024.06.18.599620

**Authors:** Enze Chen, Byumsu Kim, Nikolaos Bouklas, Lawrence J. Bonassar, Stavros Gaitanaros

## Abstract

Collagen scaffolds seeded with human chondrocytes have shown great potential for cartilage repair and regeneration. However, these porous scaffolds buckle under low compressive forces, creating regions of highly localized deformations that can cause cell death and deteriorate the integrity of the engineered tissue. We perform three-dimensional (3D) tomography-based characterization to track the evolution of collagen scaffolds’ microstructure under large deformation. The results illustrate how instabilities produce a spatially varying compaction across the specimens, with more pronounced collapse near the free boundaries. We discover that, independent of differences in pore-size distributions, all collagen scaffolds examined displayed strong auxetic behavior i.e., their transverse area contracts under compression, as a result of the instability cascade. This feature, typically characteristic of engineered metamaterials, is of critical importance for the performance of collagen scaffolds in tissue engineering, especially regarding the persistent challenge of lateral integration in cartilage constructs.

## Introduction

Over the past three decades, tissue-engineered cartilage constructs have exhibited promising potential in restoring native tissue functionality (1). Currently available, or at a late-stage clinical trial, cartilage products (such as MACI^@^, NeoCart^@^, and CMI) employ soft porous materials, such as collagen foams, to serve as scaffolding microstructures that provide initial attachment sites and mechanical support for chondrocytes (2–4). These scaffolds are designed to contain vertical pores to promote the generation of a matrix that mimics the oriented microstructure of native cartilage. The resulting mechanical properties of these collagen scaffolds are of significant importance in these biofabrication approaches, given that the tissue-engineered products undergo in vivo compression following implantation. Additionally, the Food and Drug Administration (FDA) guidance suggests that the mechanical characterization of engineered cartilage products is important in understanding implant performance (5). Despite the known importance of the underlying mechanics, no tissue-engineered cartilage products to this date have been able to recapitulate all mechanical properties of native tissue (6). Previous studies have highlighted the significant impact of morphological features of tissue-engineered cartilage on its micromechanical behavior and failure modes in the form of local strain concentrations (7, 8). The latter have been shown to affect the viability of the cells seeded in the construct (9, 10). Collectively, these results indicate the need to quantify the effect of key microstructural features of collagen scaffolds on the resulting mechanical behavior of tissue-engineered cartilage constructs.

While previous findings have shed light on the correlation between the micromechanical environment and cell health, they primarily explored this behavior via two-dimensional (2D) analysis, where 2D imaging and sample preparation impart artifacts on material behavior (7, 8). For example, buckling that occurs out of plane with respect to the field of view cannot be captured by the experimental 2D imaging technique. Further, to enable 2D imaging, samples must be sectioned, which changes the boundary conditions compared to the intact specimen. To overcome these challenges, non-destructive 3D imaging techniques utilizing micro-computed tomography (µCT) have been used to measure 3D deformation fields of porous material in situ (11, 12). However, these techniques have not been extensively applied to understand the mechanics of soft collagen foams.

The objective of this study is to investigate the interplay between the 3D porous microstructure and the corresponding nonlinear mechanics of collagen scaffolds through µCT imaging. We focus on commercially available porous collagen scaffolds made from bovine dermal type I collagen with honeycomb and sponge architectures (13, 14) manufactured from a freeze-drying process and examine their compressive behavior. In particular, we first perform microstructure characterization and use image analysis to extract key morphological features for both honeycomb and sponge scaffolds. In-situ compressive experiments with sequential µCT scans are used to highlight deformation patterns, identify the corresponding critical buckling modes, and quantify the evolution of key microstructural features at different levels of macroscopic strain. Further, the µCT technique enables us to identify the microscopic mechanisms through which an emerging macroscopic auxetic response manifests. Establishing this auxetic response enables the identification of novel micromechanical mechanisms for implant failure in vivo. The reported data are vital for inverse-engineering cartilage constructs with tailored behavior at the post-buckling regime for optimal tissue regeneration.

## Results

### Microstructure characterization of collagen scaffolds

We first focus on quantifying key morphological features of collagen constructs with honeycomb and sponge microstructures. Figure 1(a) and Figure 1(b) show the reconstructed solid models extracted from the µCT scans. Although both scaffolds share a similar quasi-2D tubular microstructure, the associated pores have distinct characteristics. Using 2D section images from approximately the mid-height of each specimen, we extract the distributions of four structural descriptors that have been shown (15–17) to greatly influence the resulting mechanical properties of porous materials: (i) wall thickness, (ii) pore area, (iii) pore compactness, and (iv) neighbor distance. Figure 1(c) illustrates the mean wall thickness for each scaffold across the specimen height. The results reveal that there are no significant density gradients across each specimen, with the wall thickness for both scaffolds being nearly constant throughout their domain. The honeycomb construct is shown to have a slightly higher average wall thickness (6.42µm) than the sponge scaffold (6.34µm). Dividing the height of each specimen (∼1.2mm) with the corresponding mean wall thickness gives a slenderness ratio of 175 and 189 for the honeycomb and the sponge respectively. These values confirm that under compression, elastic buckling will be the governing deformation mechanism. The distribution of pore areas for the two collagen constructs is displayed in Figure 1(d). The honeycomb scaffold exhibits two notable peaks at approximately 39520µm^2^ and 16750µm^2^. On the contrary, there is a significantly larger number of smaller pores within the sponge scaffolds, resulting in a single peak of the distribution at 5025µm^2^ pore area. Despite the differences in pore-size distributions, the shape of the pores in both structures is similar, as indicated by the compactness metric distribution shown in Figure 1(e). The honeycomb demonstrates an average compactness of 0.68±0.16, that is slightly higher than the corresponding values for the sponge (0.66±0.14). These values indicate that most of the pores in the collagen scaffolds are not quite circular, since their compactness is closer to the one of a square i.e., *π*/4. Finally, in Figure 1(f) we report the pore neighbor distance distribution for both types of scaffolds. The honeycomb displays a smoother distribution than the sponge scaffold due to the larger and more uniform size of its pores. Furthermore, in both honeycomb and sponge scaffolds the neighbor distance seems to be independent of location within the sectional area, indicating homogeneous pore distributions within the structures.

**Fig. 1.**
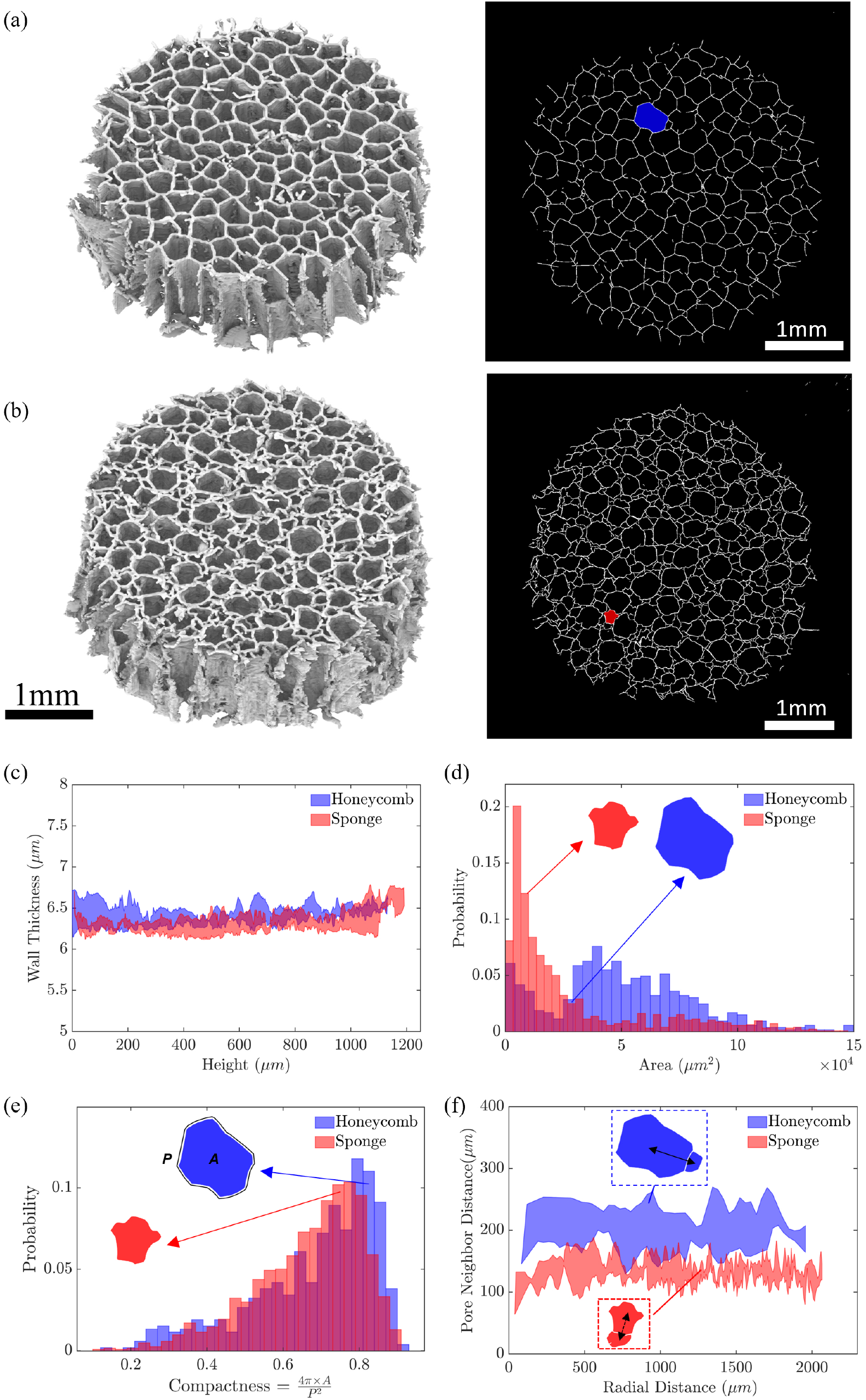
Tomography-based structure characterization of collagen constructs. (a) Reconstructed 3D solid model (left) and cross-sectional slice at mid-height (right) for a scaffold with honeycomb structure. (b) Reconstructed 3D solid model (left) and cross-sectional slice at mid-height (right) for a collagen scaffold with sponge structure. (c) Mean wall thickness across the specimens’ heights. (d) Pore area distributions. (e) Pore compactness distributions. (f) Neighbor distance as a function of radial distance.

### In-situ Testing and Deformation

Figure 2(a) illustrates the evolution of collapse for a collagen scaffold with honeycomb structure at six levels of applied macroscopic strain. To capture the complex and multi-scale nature of the involved deformations, we extract images of the 3D reconstructed specimens (top), as well as 2D sections across all three directions (bottom) for all increments of the compressive loads. Initially, the walls of the undeformed scaffold remain nearly straight, though at an inclined angle with respect to the compressive direction. In step 2, with an applied strain close to 8.5%, the walls of certain cells near the boundary start buckling into the scaffold, as observed from the 1-1 and 2-2 sections (see circled regions). Contrary, most cell walls within the structure retain their initial orientation. With increasing compression (22.5%), the collapse of pores propagates towards the interior of the cellular microstructure. These deformations at the cell level cause a notable reduction in the cross-sectional area of the whole specimen, as clearly evident from the corresponding top view. As the compaction of pores grows significantly (*>*30%), contact between neighboring walls provides additional support and prevents the further propagation of collapse near the center of the scaffold.

**Fig. 2.**
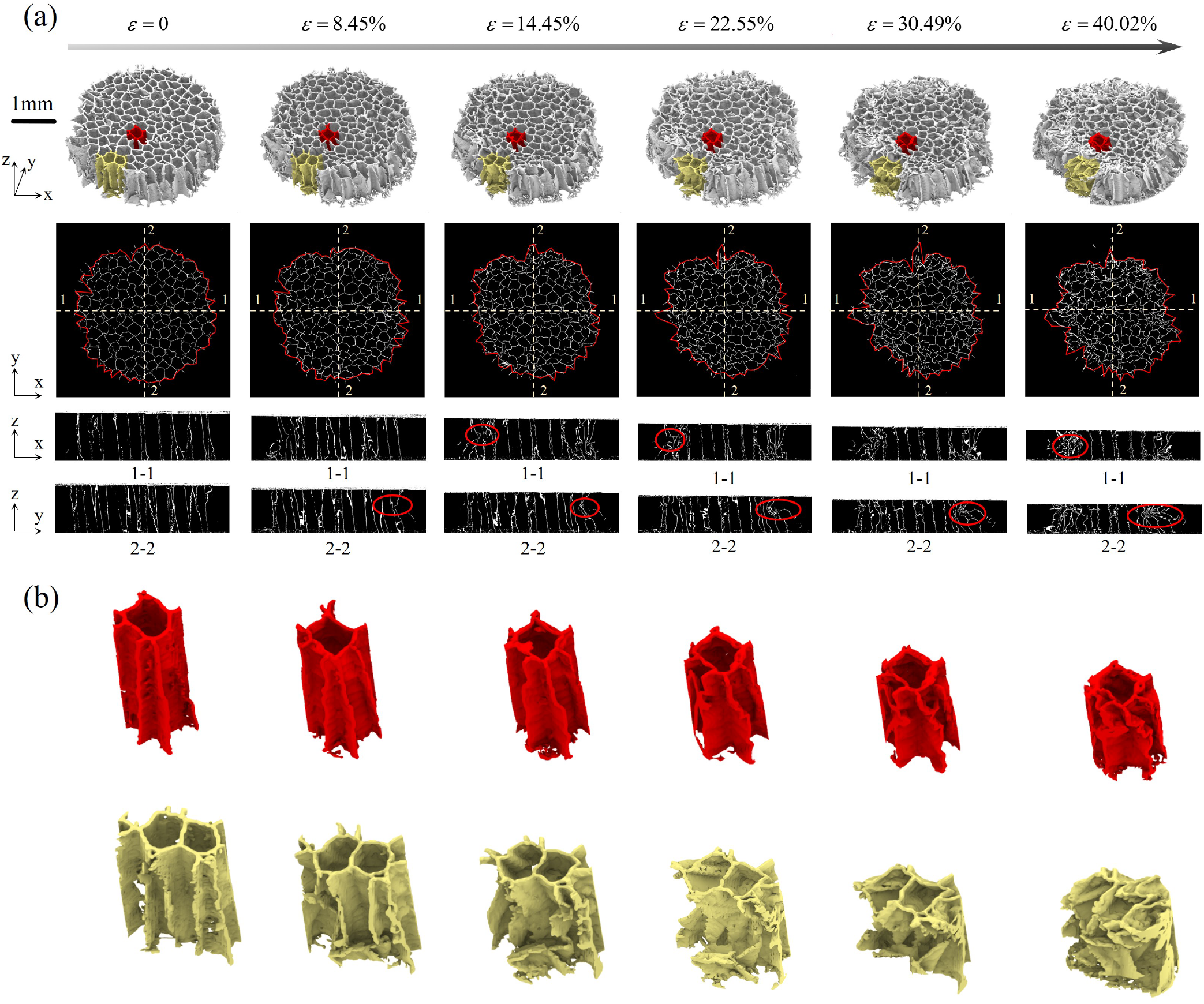
In-situ buckling and localization in honeycomb construct. (a) Imaging of large deformation evolution through (top to bottom): reconstructed 3D solid models, cross-sectional slices at mid-height, and transverse slices at six levels of applied macroscopic strain. (b) Buckling of cell walls and deformation of pores near the center (red) and the boundary (yellow).

To examine the local deformation of scaffold walls and their spatial dependence, we extract two clusters of cells, one near the center (marked with red) and one close to the boundary (marked with yellow) and monitor their individual structural evolution during compression (Figure 2(b)). It is seen that the central pores mainly maintain their overall shape, even for large macroscopic strains, depicting a uniform manner of collapse through the wrinkling of the scaffold walls. In contrast, the boundary cells show significant local and global deformations at similar levels of compression, which in turn result in increased pore compaction.

We subsequently repeat the in-situ testing and monitoring process for scaffolds with a sponge microstructure (Figure 3). It is again observed that the thin walls of the scaffold are initially straight, however in this case they also appear to be nearly vertical i.e., displaying a less pronounced inclination with respect to the loading axis than the walls of the honeycomb scaffold. At imaging level 3, corresponding to a macroscopic strain of approximately 30%, we clearly observe the buckling of boundary cells in the 1-1 and 2-2 sections. Interestingly, the deformation of the cell walls varies in the vertical direction too, with the appearance of walls that have localized near the bottom, as well as ones that collapsed at mid-height. As compression progresses, the bottom region continues to collapse due to buckling, causing a shift of the scaffold, as evidenced in the 1-1 section. Imaging in the 2-2 sections reveals an inward-type of folding that again results in an overall decrease of the specimen cross-sectional area. Figure 3(b) illustrates the corresponding evolution of pore morphology for clusters close to the center (top, red) and boundary (bottom, yellow). In a similar manner to the honeycomb construct, the pores located closer to the center maintain their shape to a greater extent than those near the boundary that experience significant compaction and appear nearly densified at 47% strain.

**Fig. 3.**
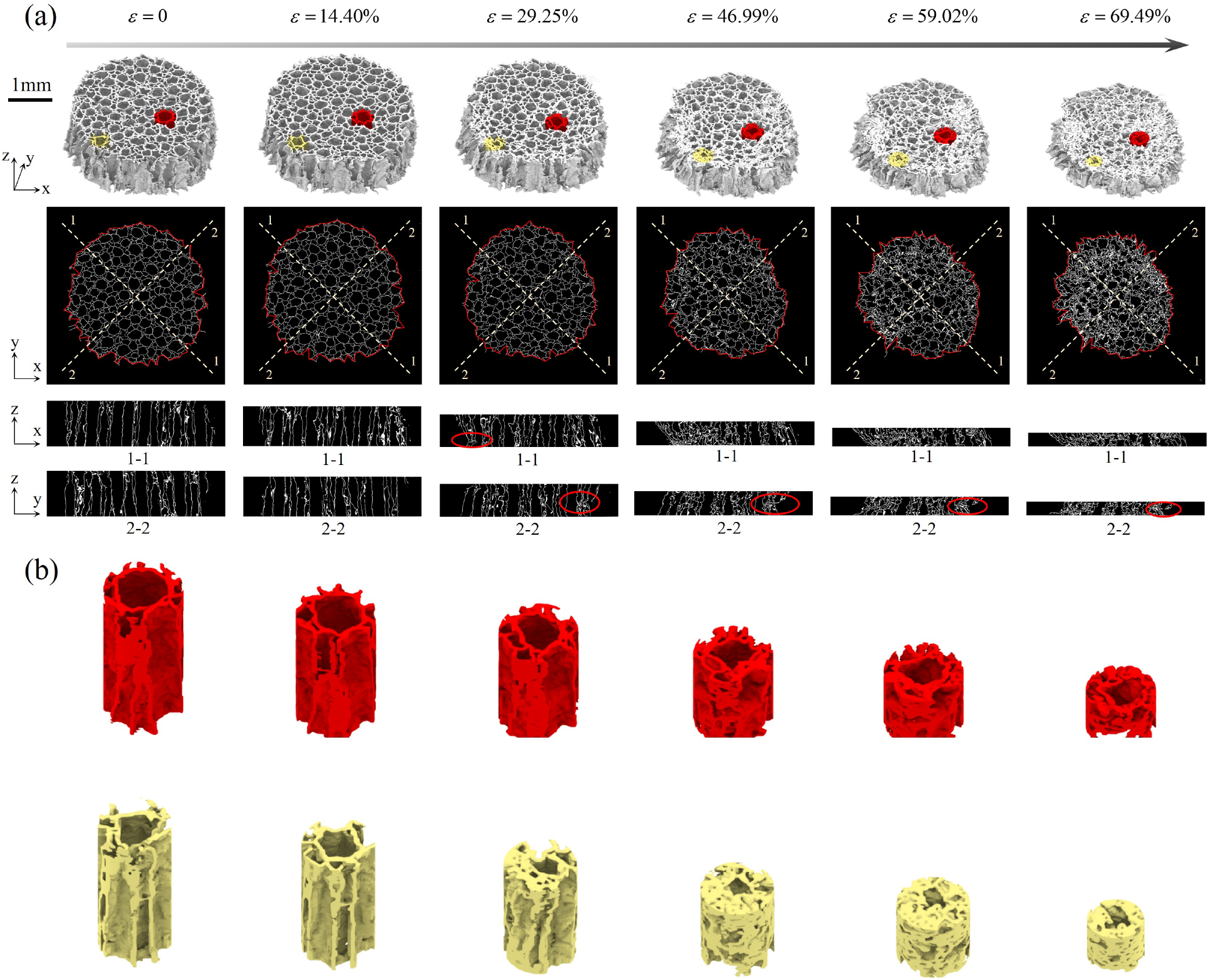
In-situ buckling and localization in sponge construct. (a) Imaging of large deformation evolution through (top to bottom): reconstructed 3D solid models, cross-sectional slices at mid-height, and transverse slices at six levels of applied macroscopic strain. (b) Buckling of cell walls and deformation of pores near the center (red) and the boundary (yellow).

We further exploit the tomography imaging from the in-situ tests to extract the evolution of structure characteristics at different levels of deformation for both the honeycomb Figure 4(a-d) and sponge (Figure 4(e-h)) scaffolds. To examine the spatially varying collapse of pores, we measure the pore area (Figure 4(a/e)) and pore neighbor distance (Figure 4(b/f)) as a function of radial distance from the center. In both cases, the corresponding distributions for the undeformed and highly deformed scaffold are compared. It is evident that the collapse of pores results in a significant reduction in both morphological descriptors, regardless of the particular scaffold morphology. Furthermore, this decrease is more pronounced at distances larger than 1mm from the center of the specimens. These findings provide quantitative evidence of the observations made based on the 3D reconstructed images shown in Figure 2(a) and Figure 3(a). Figure 4(c) and Figure 4(c) depict the specimens’ cross-sectional areas, corresponding to the domain enclosed by the red curve in Figure 2(a)-Figure 3(a), as a function of height, for six increments of the applied strain. A key observation that is common for both microstructures involve the decrease of macroscopic area with increasing deformation. However, there is a notable difference between them. The honeycomb structure displays a non-uniform cross-sectional area across its height, characterized by a clear dip in the middle. This variation arises from buckling occurring at the structure’s midheight, leading to a significant area reduction in these regions. Conversely, the sponge structure maintains a more consistent cross-sectional area distribution throughout its height but has the lowest area near the bottom boundary. The sponge’s even distribution is attributed to its buckling location near the base and slanted shape in post-buckling, which concentrates the buckling at its lower boundary. This results in a uniform area reduction in the other regions. As a result, the distribution of the cross-sectional area highly correlates with the buckling location and post-buckling behavior and can be used to probe the deformation of the scaffold.

**Fig. 4.**
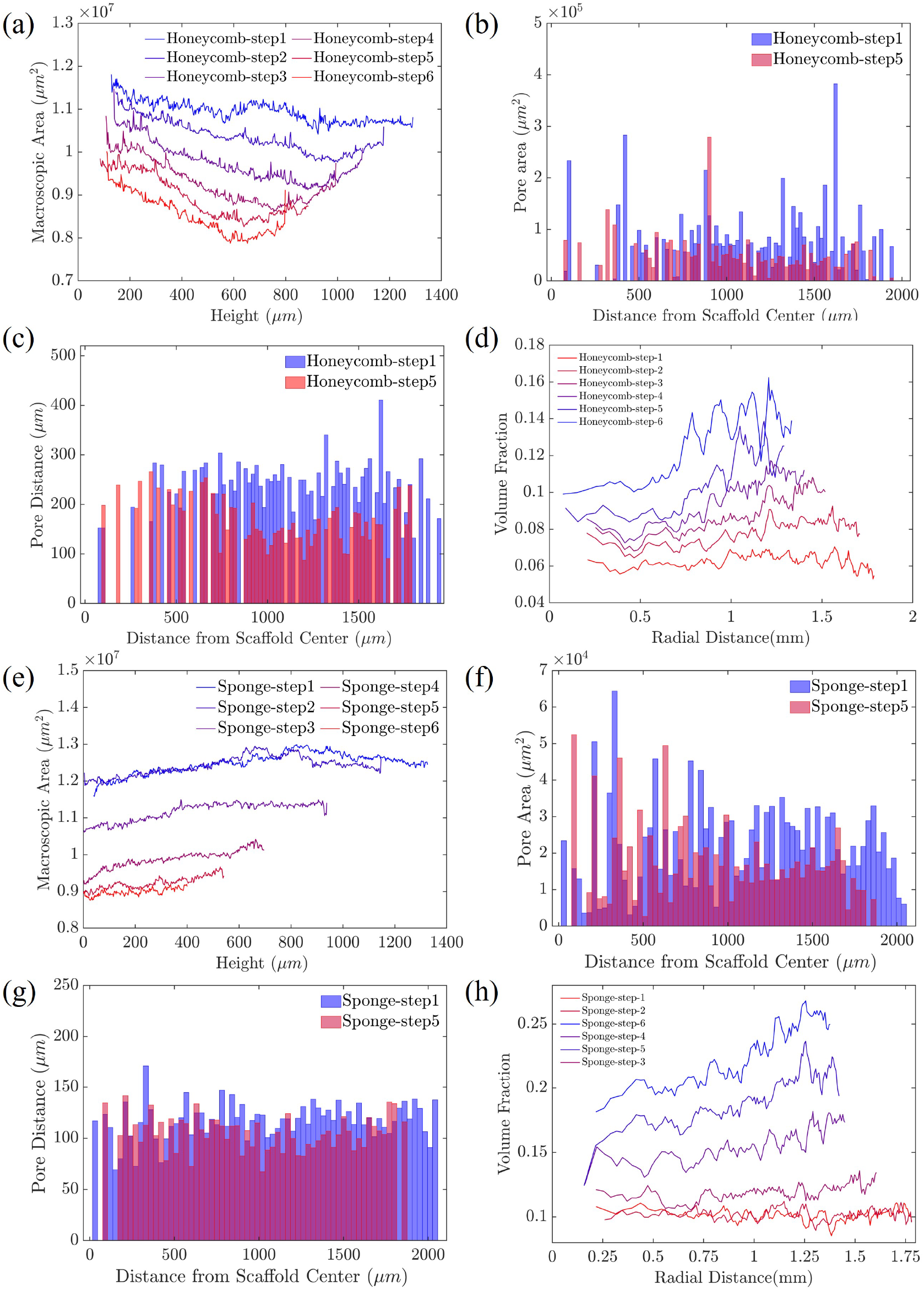
Evolution of microstructural features of a honeycomb construct with increasing deformation. (a) Macroscopic area as a function of specimen height. (b) Pore area distribution as a function of radial distance in the undeformed (ϵ = 0) and deformed (ϵ = 30.5%) configurations. (c) Neighbor distance as a function of radial coordinate in the undeformed (ϵ = 0) and deformed (ϵ = 30.5%) configurations. (d) Spatial distribution of volume fraction in different strain levels. Evolution of microstructural features of a sponge construct with increasing deformation. (e) Macroscopic area as a function of specimen height. (f) Pore area distribution as a function of radial distance in the undeformed (ϵ = 60%) and deformed (ϵ = 0) configurations. (g) Neighbor distance as a function of radial coordinate in the undeformed (ϵ = 0) and deformed (ϵ = 60%) configurations. (h) Spatial distribution of volume fraction in different strain levels.

To further examine the compaction of the scaffold under compression, we analyzed the volume fraction in various regions based on the radial distance for both honeycomb and sponge, as shown in Figure 4(d) and Figure 4(h). Both structures share one key observation of heterogeneous volume compaction in the radial direction. Initially, both structures exhibit nearly uniform volume fractions radially. However, as compression continues, the volume fraction increases more significantly farther from the center. Specifically, regions beyond a radial distance of 700µm display a pronounced jump in volume fractions when highly compressed. In the honeycomb structure, regions up to 700µm from the center see an average volume fraction increase of 74% from its original state to a 40% compressed state. In contrast, areas beyond 700µm experience a much higher volume fraction increase of 122%. For sponges, the central regions only exhibit an average 91% growth in volume fraction, while the volume fraction shows a 133% increase in the outer regions. This finding quantitatively confirms the scaffold’s non-uniform compaction and concentrated deformation caused by the instability at the boundary and the post-buckling behavior observed in Figure 2 and Figure 3.

Nonetheless, the significant reduction of specimen area with increasing compaction seems to be a feature that is independent of the specific scaffold morphology. To highlight this feature, we plot for all collagen scaffolds tested, their cross-sectional area at mid-height A, normalized by its undeformed value *A*_0_, as a function of the applied strain (see Figure 5(a)). It is obvious that all specimens see a decrease that for a 40% strain varies between 5-25%, This result indicates a strong auxetic behavior of collagen constructs, that is very different than typical monolithic solids that expand in the direction normal to the compressive loads. To illustrate the striking difference between a typical soft material and these collagen constructs we include in Figure 5(a) the corresponding area evolution of a Neo-Hookean material under compression. The associated deformations for a 40% nominal strain are shown in Figure 5(b). Even though many soft cellular materials, including elastomeric foams (18) and 3D-printed porous media (19), show negligible expansion in the transverse direction when compressed, the strong area reduction that the collagen scaffolds display, corresponding to a negative tangent Poisson’s ratio, is a characteristic seen in a certain class of mechanical metamaterials (20–22) and is often associated with improved energy absorption characteristics.

**Fig. 5.**
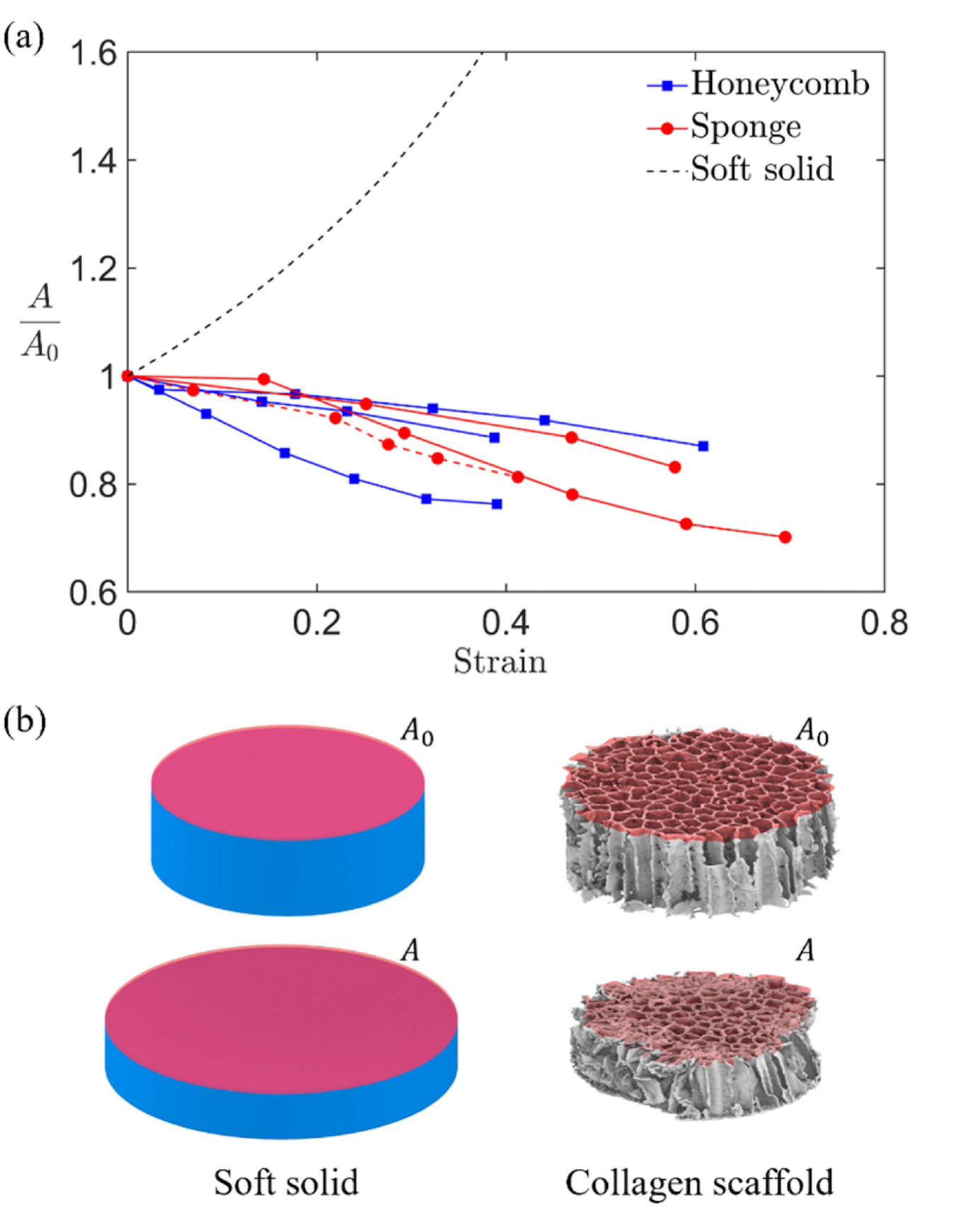
(a) Evolution of normalized macroscopic area with increasing macroscopic strain for collagen scaffolds with honeycomb and sponge microstructures. For comparison, the corresponding area evolution of a monolithic soft solid is included. (b) The gradual reduction in the cross-sectional area shows the strong auxetic metamaterial behavior of collagen constructs i.e., the presence of lateral contraction when vertically compressed, instead of the expansion observed in typical soft solids.

## Discussion

In summary, here we present a complete 3D microstructural analysis of soft collagen scaffolds with honeycomb and sponge microstructures under compressive loads. We find that the resulting deformation is highly heterogeneous across the specimen area with the emergence of regions with increased pore compaction driven by elastic instabilities. This compaction is attributed to the microstructural design of collagen scaffolds and the vertically oriented pores with a length scale of ∼ 100µm. While native cartilage shares a similar vertical alignment, the associated length scales are much smaller, with collagen fiber sizes of ∼ 100nm, a feature that could alter the resulting mechanical behavior. Typically, in low-density lattices and porous materials under compression, localization leads to bands of collapsed pores forming at a certain slope with respect to the loading surface (18). Contrary, in this work the collapsing pores form peripherally and divide the scaffold in highly deformed pore clusters near the free boundary, and much less distorted regions in the interior of the specimen. This behavior is reminiscent of the presence of floppy modes on the free boundaries of lattice structures (23). The increased compaction of the collapsed pores near the specimen lateral boundary, and the associated contact between neighboring walls could be responsible for obstructing the propagation of buckling towards the interior of the scaffold. More importantly, the cross-sectional area of the collagen scaffolds decreases gradually as compression increases and the collapse of pores progresses. This area reduction is attributed to the inward buckling and folding of the pore walls. At large deformations, corresponding to effective macroscopic strains of 40%, the scaffolds’ area at their mid-section is reduced by 5-25%. This auxetic behavior, i.e. the displayed contraction/elongation of the material in the transverse direction when compressed/stretched in the longitudinal one, has been reported extensively in cellular solids and in particular those with re-entrant members (24) or chiral mesostructures (25). Since in these materials the resulting negative Poisson’s ratio is driven by geometry, there have been efforts (26–28) to exploit these microstructural features in additively manufactured scaffolds for tissue engineering. In contrast to these material systems, the auxetic behavior in the collagen scaffolds examined here is deformation-dependent and generated by the elastic instabilities that govern the associated large deformations. Even though these instabilities can potentially lead to beneficial properties (20–22), in the case of collagen scaffolds in cartilage constructs the auxetic behavior may well impede implant integration. To date, the lateral integration of engineered cartilage constructs with host tissue has proven to be a consistent challenge (29). Surprisingly, past in vivo studies have demonstrated that the lateral integration of tissue-engineered cartilage constructs using porous collagen scaffolds is not significantly superior to that achieved through microfracture surgeries (30–32). In microfracture surgeries, insufficient lateral integration often results from the disparity in cartilage types between the newly formed fibrocartilage and the native host hyaline cartilage, or incomplete defect fill (33–37). Conversely, tissue-engineered cartilage constructs created from porous scaffolds do not encounter these issues, as they are composed of hyaline cartilage and fully occupy the defect. Therefore, the fundamental reason for the persistent challenge in lateral integration of implanted tissue-engineered cartilage constructs remains poorly understood. The lateral contraction of the collagen scaffolds reported here due to auxetic behavior, could result in the formation of a physical gap between the engineered cartilage constructs and the native host tissue. In addition, the increased compaction near the boundary pores can induce cell death (10), resulting in significantly lower cell viability at the vicinity of the free boundary compared to the interior of the engineered construct. Previous studies have indicated that the low cell viability at the interface between the graft and the host tissue can hinder the integration process (29, 38, 39). Overall, the auxetic behavior observed in collagen scaffolds provides valuable insights into the underlying reasons for the persistent challenge of lateral integration in tissue-engineered cartilage constructs. It is important to note that the experiments reported here do not aim to replicate the behavior of the complete cartilage constructs in vivo. Doing so would require a different testing setup as well as filling partially the pores of the scaffolds with chondrocytes and matrix, which are left for future studies. Nonetheless, understanding the effect of microstructure on the resulting elastic instability cascade and evolution of collapse, in an uncoupled manner from the complex micromechanical environment during implantation is deemed imperative. Furthermore, high-fidelity numerical models that are able to reproduce experimental data and facilitate the exploration of the vast parameter space are also essential for designing novel microstructures. Towards this goal, advances in 3D bio-printing are expected to provide increased control over the structure of synthesized scaffolds, and potentially enable the tailoring of morphological features that yield pre-determined target deformation modes at the post-buckling regime.

## Materials and Methods

### Collagen scaffold preparation

Honeycomb (Histogenics Corp., Waltham, MA) and sponge (Koken CO., LTD, Tokyo, Japan) collagen scaffolds were obtained. Both honeycomb and sponge scaffolds were made from type I bovine dermis collagen and had pore sizes ranging from 100 - 200µm in diameter and 1.5mm in height, according to manufacturing specifications. A total of 6 samples (3 honeycomb scaffolds and 3 sponge scaffolds) were cut using 4mm and 6mm biopsy punches (Integra York PA, Inc., York, PA) with pores aligned in the axial direction.

### Micro-computed tomography

The cellular microstructure of collagen constructs is characterized through µCT using a Skyscan 1172 system (by Bruker). The scanning and imaging settings adopted, based on maximizing accuracy and performance, include 67kV and 174µA power for the X-ray source, projection images over a 180° rotation without any filtering, 2.3µm image pixel size. The projection images underwent flat-field and dark-field corrections for better contrast. After scanning, projection images are reconstructed by the NRecon software with default post-alignment, smoothing, and ring artifact correction settings.

### Image analysis and extraction of microstructural characteristics

2D slices obtained from the reconstructed images are imported into MATLAB for further imaging analysis and extraction of key morphological characteristics. We employ the hysteresis thresholding method to differentiate between collagen and void regions. Following this approach, pixels with intensity values above a user-set upper threshold are classified as solid, while pixels with intensity values below a corresponding low threshold are classified as voids. All pixels with intermediate intensity values are then evaluated based on their connectivity to the pixels corresponding to the solid phase. Here, a pixel connectivity parameter equal to four is chosen. We extract next the distributions of four microstructural characteristics, namely the wall thickness, pore area, compactness, and pore neighbor distance along the height of the scaffold, at four equidistant increments. The wall thickness t around each pore is determined by the bwdist function in MATLAB. The size and shape of each pore are characterized by calculating the corresponding area A and compactness. The latter is estimated using a ratio of the pore area A and the pore perimeter P i.e., 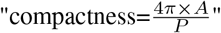. The pore “density” is evaluated through the neighbor distance metric i.e., the length between the centroids of two adjacent pores. All of the above calculations require distinct boundaries for all pores, that we achieve here by skeletonizing the binary 2D images using the bwboundaries function in MATLAB. For the 3D visualization and image analysis, each binary image stack is exported to ImageJ where, using the 3D viewer function with a resampling factor of two, a surface mesh is generated. The mesh is then exported as an STL file into Rhino3D for visualization.

### In-situ testing

We conduct a series of in-situ compressive experiments under displacement control using an MTS loading stage with a 44N load cell. In each test, the specimen is placed in the center of the stage and subsequently an upward displacement with a rate (∼ 0.5mm/min) is prescribed on the bottom platen to ensure quasi-static compaction. Once the specimen comes to full initial contact with the platen, the scaffold is scanned and its undeformed microstructure is extracted. Subsequently each specimen is compressed in increments of average macroscopic strain of 7-9%. At the end of each loading increment, testing is paused, and the specimen is scanned, keeping the image acquisition settings consistent. These steps are iterated until each specimen reaches an average macroscopic compaction of 40-70%. We use here a set of three samples for each cellular microstructure i.e., for the honeycomb and sponge constructs.

### Deformation-dependent morphological features

Through the in-situ testing of the collagen scaffolds, we further calculate the evolution of morphological characteristics as a function of the applied loading and resulting deformation. In addition to the aforementioned pore-related features, here we also measure how increased compression affects two additional microstructure properties: (a) the sectional area of each specimen across their height and (b) the volume fraction as a function of radial distance from the specimen center. Regarding the former, we first calculate the sectional area centroid of the scaffold by averaging the binary image with its position, and then create 100 circumferential curves from the center to the image boundary. Subsequently, we calculate the outer intersection point between each curve and the scaffold. We connect all these intersection points in a right-handed order to form a closed domain and measure the corresponding area. For the latter part, the spatial distribution of the volume fraction was quantitatively assessed using a sliding box approach on binary image stacks of the scaffold from the scanning. A three-dimensional box of a predefined x-y size (460µm) was systematically slid across this binary domain in the x-y plane with a specific stride (230µm), spanning the entire depth of the stack in the z-dimension to ensure comprehensive assessment across the scaffold’s height. For each box, the volume fraction was determined as the ratio of voxels of the scaffold to the total number of voxels in the box. Simultaneously, the Euclidean distance between the center of each box and the scaffold’s central point was calculated. This approach enabled a quantitative analysis of the heterogeneous compaction of the scaffold during the compression.

## ACKNOWLEDGEMENTS

The authors would like to thank Robert C. Spiro (Aesculap Biologics, LLC) for helpful discussions.

## Author contributions

Conceptualization: NB, LJB, SG Investigation: EC, BK Supervision: NB, LJB, SG Funding acquisition: NB, LJB, SG Writing—original draft: EC, BK Writing—review & editing: EC, BK, NB, LJB, SG

## Data and materials availability

All data needed to evaluate the conclusions in the paper are present in the paper and/or the Supplementary Materials.

